# N-Methyl-D-aspartate receptor hypofunction causes recurrent and transient failures of perceptual inference

**DOI:** 10.1101/2024.05.24.595590

**Authors:** Veith Weilnhammer, Marcus Rothkirch, Deniz Yilmaz, Merve Fritsch, Lena Esther Ptasczynski, Katrin Reichenbach, Lukas Rödiger, Philip Corlett, Philipp Sterzer

**Affiliations:** Department of Psychiatry, Charité-Universitätsmedizin Berlin, corporate member of Freie Universität Berlin and Humboldt-Universität zu Berlin, Germany; Berlin Institute of Health, Charité-Universitätsmedizin Berlin and Max Delbrück Center, Germany; Helen Wills Neuroscience Institute, University of California Berkeley, USA; Medical School Berlin, Hochschule für Gesundheit und Medizin, Germany; Berlin School of Mind and Brain, Humboldt-Universität zu Berlin, Germany; Max Planck School of Cognition, Leipzig, Germany; Department of Psychiatry and Psychotherapy, LMU University Hospital, Munich, Germany; Department of Psychiatry, Yale University School of Medicine, New Haven, USA; Department of Psychiatry (UPK), University of Basel, Switzerland

## Abstract

Perception alternates between an external mode, driven by sensory inputs, and an internal mode, shaped by prior knowledge. We found that the external mode is more prevalent during pharmacologically induced N-Methyl-D-aspartate receptor (NMDAR) hypofunction and in schizophrenia. This NMDAR-dependent increase in external mode suggests that psychotic experiences are caused by recurring dissociations of perception from prior knowledge about the world.

## 2 Main

Imagine a dimly lit room at a crowded party, where unclear visual signals, indistinct sounds, and complex social interactions allow for multiple - and sometimes false - interpretations. In such ambiguity, failures of perceptual inference, the ability to contextualize sensory inputs with prior knowledge about the world, can lead to profound departures from reality: Faces obscured in shadow may appear distorted, random noise could be perceived as a whisper, and friendly smiles might seem derogatory.

This example illustrates why a disruption of perceptual inference is likely to play a crucial role in schizophrenia (Scz), a chronic and severe mental disorder characterized by psychotic symptoms such as delusions and hallucinations^1–3^. Yet despite considerable progress in the computational understanding of psychosis, two key questions have remained unanswered.

The first question concerns the neural mechanisms that cause perceptual inference to fail in Scz. Several lines of evidence point to N-Methyl-D-aspartate receptor (NMDAR) hypofunction as a key factor in the pathophysiology of psychosis^4^. NMDAR antibodies^5^ and antagonists such as ketamine^6^ mimic the symptoms of Scz, which is itself associated with a reduction of NMDAR density in prefrontal cortex^7^. NMDARs control the ratio of neural excitation and inhibition^8^, block the release of midbrain dopamine^9^, enable cortical feedback^10^ and support synaptic short term plasticity^11^. While these NMDAR-dependent mechanisms are likely critical for perceptual inference, it is yet to be determined how NMDAR hypofunction may cause the psychotic symptoms that characterize Scz.

The second unresolved question concerns the temporal dynamics of psychotic experiences, which often unfold as short-lived events spanning from seconds to minutes, especially at early stages of Scz^12,13^. The transient nature of psychotic experiences challenges models that assume a constant disruption of perceptual inference^1–3^. A solution to this problem is suggested by the recent observation that perceptual inference is subject to spontaneous fluctuations over time that occur at a timescale compatible with the duration of individual psychotic experiences^14–16^. Such fluctuations have been related to two opposing modes of inference, during which perception is driven predominantly either by external inputs or by internal predictions that stem from recent perceptual experiences^17^ (Figure 1A). Although preliminary evidence indicates a tendency toward the external mode in people with Scz^18^, the neural mechanisms of mode fluctuations and their potential implications in psychosis have remained elusive.

**Figure 1.**
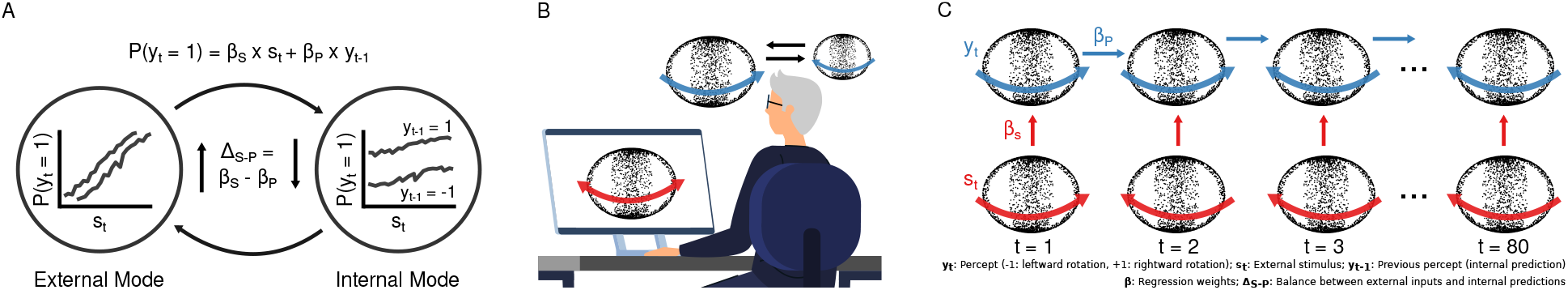
A. When inferring whether the world is one state or another (*P* (*y*_*t*_ = 1) or *P* (*y*_*t*_ = 0), respectively), the brain integrates ambiguous sensory signals *s*_*t*_ with internal predictions that reflect prior knowledge about the environment. One source of prior knowledge is the temporal autocorrelation of natural stimuli, in which the recent past often predicts the near future. The integration of external inputs and internal predictions depends on the weights assigned to incoming sensory data (*β*_*S*_ *× s*_*t*_) and to internal prediction derived from previous experiences (*β*_*P*_ *× y*_*t−*1_, dotted versus solid lines, simulated data), respectively. *β*_*S*_ determines the slope, and *β*_*P*_ the shift of the psychometric function that links *s*_*t*_ and *y*_*t*_. Importantly, the balance Δ_*S−P*_ = *β*_*S*_ *− β*_*P*_ is known to alternate between two opposing modes: During the external mode (left), perception is largely determined by *β*_*S*_ *× s*_*t*_, which is reflected by a steep slope and a small shift of the psychometric curve. Conversely, during the internal mode (right), perception is shaped by *β*_*P*_ *× y*_*t−*1_, resulting in a shallow slope and a large shift of the psychometric curve. B. We conducted a double-blind placebo-controlled experiments in 28 healthy human participants, who received a continuous infusion with either the NMDAR antagonist S-ketamine or saline. During the infusion, the participants viewed an ambiguous SFM stimulus that was compatible with two mutually exclusive subjective experiences (left vs. rightward rotation of the front surface, red arrows). This induced the phenomenon of bistable perception: While the ambiguous stimulus remained constant, participants perceived only one direction of rotation (blue arrow), before switching to the competing alternative. C. Changes in the perceived direction of rotation of the SFM stimulus occur at brief depth-symmetric configurations of the stimulus (Supplemental Video S1). We therefore transformed the behavioral responses into a sequence of states *t* (80 1.5 sec intervals per block), each associated with a combination of the SAR-weighted input *s*_*t*_ and the perceived direction of rotation *y*_*t*_. We used GLMs to quantify the weights *β*_*S*_, *β*_*P*_ and *β*_*B*_, which reflect how inferences *y*_*t*_ were determined by the external inputs *β*_*S*_ *× s*_*t*_ and internal predictions *β*_*P*_ *× y*_*t−*1_.

The objective of the current study was therefore twofold: First, to test whether NMDAR hypofunction causes changes in perceptual inference that characterize Scz; and second, to explore the effect of NMDAR hypofunction on ongoing fluctuations in perceptual inference that may explain the transient nature of psychotic experiences. We addressed these aims in a double-blind placebo-controlled cross-over experiment in 28 healthy human participants. The participants attended two experimental sessions during which they received a continuous intravenous infusion of either the NMDAR antagonist S-ketamine at a dose of 0.1 mg/kg/h or a saline placebo. In each session, the participants viewed ten 120 sec blocks of an ambiguous structure-from-motion (SFM) stimulus that induced the experience of a sphere rotating around a vertical axis, and reported changes in the perceived direction of rotation (leftward vs. rightward movement of the front surface) as well as their confidence in the choice (Figure 1B and Supplemental Video S1).

The ambiguity of the display induced the phenomenon of bistable perception, where spontaneous changes in the perceived direction of rotation occurred in average intervals of 13.75 ± 3.09 sec. In line with previous results^19,20^, these changes in perception occurred with a probability of 0.11 ± 8.67 *×* 10^−3^ at brief depth-symmetric configurations of the stimulus (see Supplemental Video S1 and Supplemental Figure S1A). We therefore divided the continuous behavioral reports into a sequence of discrete states *t*. Each state was associated with a perceptual experience *y*_*t*_, confidence *c*_*t*_ and the external input *s*_*t*_.

Bistable perception can be conceptualized as an inferential process about the cause of *s*_*t*_, in which previous experiences (*y*_*t−*1_) reflect internal predictions that provide prior knowledge about the interpretation *y*_*t*_ of the ambiguous stimulus^20^ (Figure 1C). To test how NMDAR antagonism altered the balance between external inputs and internal predictions, we attached a 3D signal to a fraction of the stimulus dots. The signal-to-ambiguity ratio (SAR) ranged from complete ambiguity to full disambiguation across five levels and remained constant in each block of the experiment. By changing the direction of rotation enforced by the 3D signal at random in average intervals of 10 sec, we created dynamic conflicts between the SAR-weighted input *s*_*t*_ and the stabilizing internal prediction *y*_*t−*1_. As expected, we found that *y*_*t*_ was driven by both *s*_*t*_ (*β*_*S*_ = 3.01 ± 0.06) and *y*_*t−*1_ (*β*_*P*_ = 2.06 ± 0.03). Importantly, S-ketamine caused perception to shift toward *s*_*t*_ (0.45 ± 0.08, z = 5.6, p = 1.71 *×* 10^−7^; Figure 2A and Supplemental Figure S2), indicating a stronger weighting of external inputs over internal predictions during pharmacologically induced NMDAR hypofunction.

**Figure 2.**
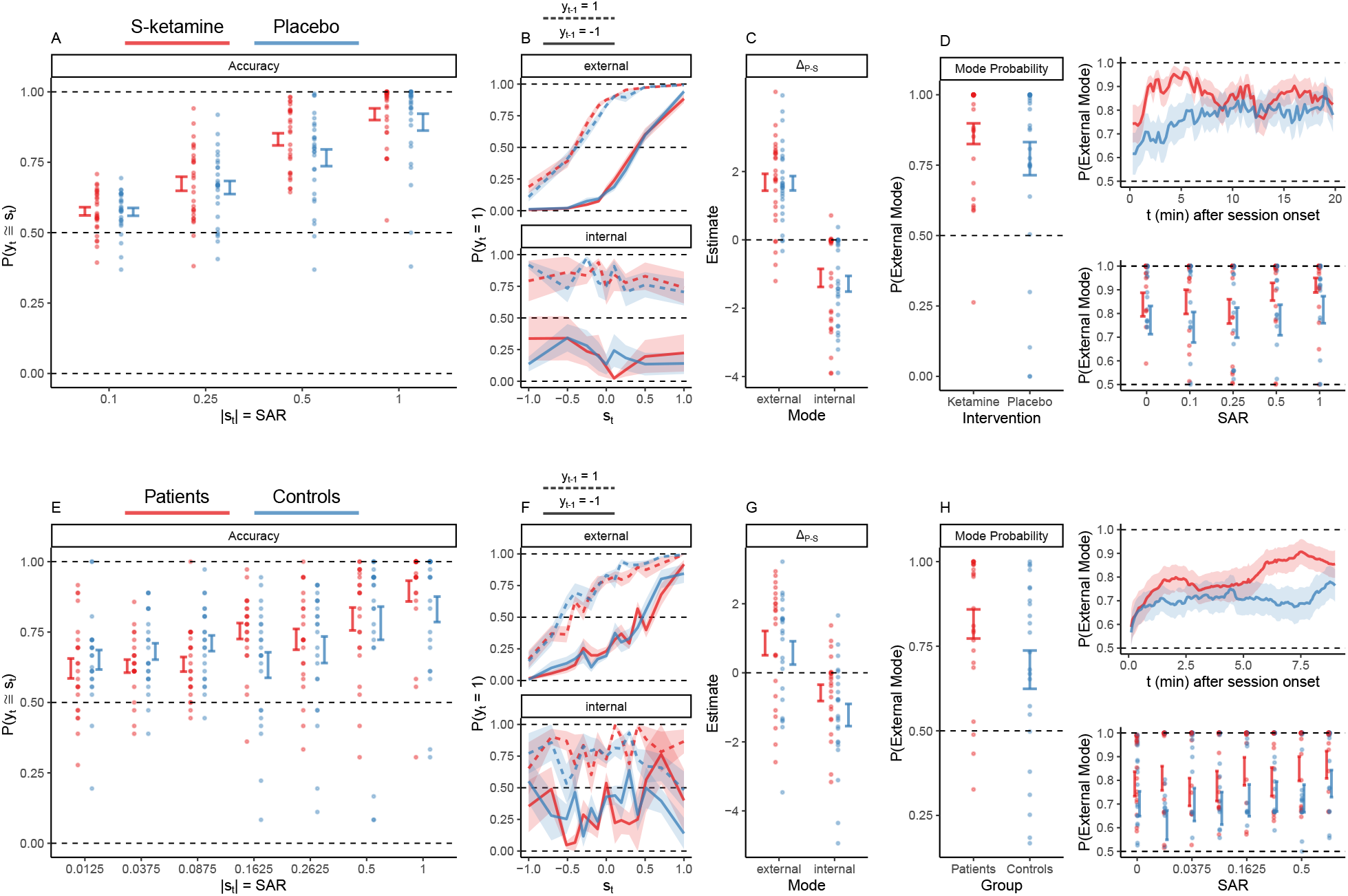
A. The percepts *y*_*t*_ were more likely to match the stimuli *s*_*t*_ at higher levels of SAR (3.01 ± 0.06, z = 50.39, p = 0). The positive effect of SAR on *P* (*y*_*t*_ ≅ *s*_*t*_) was more pronounced under S-ketamine (red) relative to placebo (blue; 0.45 ± 0.08, z = 5.6, p = 1.71 *×* 10^−7^). **B**.In the S-ketamine experiment, the HMM identified two modes that differed with respect to the relative weighting of external sensory data and internal predictions: Perception fluctuated between an external mode, determined by the input *s*_*t*_ (upper panel panel, steep slope and small shift of the psychometric curve), and an internal mode, dominated by a stabilizing prediction that biased perception toward previous experiences *y*_*t−*1_ (lower panel, shallow slope and large shift of the psychometric curve). Within modes, there was no significant effect of S-ketamine (red) versus placebo (blue) on the relation of *y*(*t*) with *s*(*t*) and *y*(*t −* 1). **C**.Δ_*S−P*_, the balance between the external input and the stabilizing internal predictions, was larger during external than during internal mode (2.8 ± 0.29, T(*−*81) = *−*9.5, p = 5.22 *×* 10^−13^). Importantly, we found no significant effect of S-ketamine (red) vs. placebo (blue) on Δ_*S−P*_ within modes (*−*0.03 ± 0.29, T(81) = *−*0.1, p = 1). **D**.S-ketamine (red) increased the probability of external mode (1.01 ± 0.03, z = 30.7, p = 4.26 *×* 10^−206^) relative to placebo (blue; left panel). The effect of S-ketamine on mode was present from the start of the session (1.77 ± 0.07, z = 26.9, p = 8.88 *×* 10^−159^, upper panel), with no significant effect of time (*−*0.18 ± 0.08, z = *−*2.17, p = 0.12). Relative to placebo, S-ketamine increased the probability of external mode across all SARs (0.85 ± 0.06, z = 14.14, p = 8.33 *×* 10^−45^, lower panel). Higher SARs were associated with an increased probability of external mode (1.34 ± 0.09, z = 15.01, p = 2.49 *×* 10^−50^), in particular under S-ketamine (0.62 ± 0.11, z = 5.52, p = 1.32 *×* 10^−7^). Alternations between external and internal mode were found at all SARs: From from full ambiguity to complete disambiguation, the probability of external mode increased by only 0.11 under S-ketamine and 0.07 under placebo. **E**.In patients (red) and controls (blue), percepts *y*_*t*_ were more likely to match the stimuli *s*_*t*_ at higher levels of SAR (*β*_*S*_ = 2.77 ± 0.11, z = 24.85, p = 2.18 *×* 10^−135^). Patients followed the external inputs more closely than controls (0.75 ± 0.15, z = 4.96, p = 5.6 *×* 10^−6^). **F**.In analogy to the S-ketamine experiment, the HMM identified two opposing modes in Scz patients (red) and controls (blue). The external mode increased the sensitivity toward *s*_*t*_ (slope of the psychometric function) and weakened the effect of the stabilizing internal prediction *y*_*t−*1_ (shift between the dotted and solid line) relative to the internal mode. Within modes, there was no effect of group on the relation of *y*(*t*) with *s*(*t*) and *y*(*t −* 1). **G**.The external mode increased Δ_*S−P*_, the balance between external inputs and internal predictions, in patients (red) and controls (blue; 1.44 ± 0.33, T(44) = 4.33, p = 3.39 *×* 10^−4^), with no significant effect of group (*−*0.28 ± 0.54, T(87.97) = *−*0.52, p = 1). **H**.Relative to controls (blue), patients (red) spent more time in external mode (0.52 ± 0.03, z = 16.88, p = 1.23 *×* 10^−63^, left panel). In both group, biases toward external mode increased over time after session onset (2.41 ± 0.11, z = *−*21.37, p = 1.02 *×* 10^−100^; upper panel), with a stronger effect in patients (1.84 ± 0.14, z = 12.97, p = 7.09 *×* 10^−38^). Patients were more likely than controls to be in external mode across all levels of SAR (0.51 ± 0.03, z = 14.56, p = 1.89 *×* 10^−47^, lower panel). External mode increased with SAR (0.63 ± 0.1, z = 6.47, p = 3.85 *×* 10^−10^), with no significant difference between groups (0.15 ± 0.13, z = 1.16, p = 0.98). As in the S-ketamine experiment, alternations between external and internal mode were found at all SARs: From from full ambiguity to complete disambiguation, the probability of external mode increased by only 0.12 in patients and 0.18 in controls.

Next, we performed the same analysis on data from a previous case-control study using an analogous task in patients with Scz^19^. In Scz patients and controls, *y*_*t*_ was influenced by the SAR-weighted input *s*_*t*_ (*β*_*S*_ = 2.77 ± 0.11) and the stabilizing prediction *y*_*t−*1_ (*β*_*P*_ = 1.5 ± 0.03). Similar to S-ketamine, *s*_*t*_ had a larger impact on perception in Scz patients than controls (0.75 ± 0.15, z = 4.96, p = 5.6 *×* 10^−6^; Figure 2E and Supplemental Figure S3). Together, these results align with the canonical predictive processing theory of Scz^1–3^: Pharmacologically-induced NMDAR hypofunction and Scz are associated with a shift of perceptual inference toward external inputs, and away from stabilizing internal predictions. NMDAR hypofunction may thus trigger psychotic experiences by causing erratic inferences about ambiguous sensory information.

As a mechanism for symptoms that are transient and recurring, NMDAR-dependent changes in perceptual inference should not be constant, but fluctuate dynamically at a timescale that is compatible with the duration of individual psychotic experiences. We tested this prediction in Hidden Markov Models (HMM) that inferred transitions between two latent states, each linked to an independent general linear model (GLM) that predicted *y*_*t*_ from *s*_*t*_ and *y*_*t−*1_. The *β* weights quantified the sensitivity to ambiguous sensory information (*β*_*S*_ *× s*_*t*_) relative to the stabilizing effect of internal predictions provided by preceding experiences (*β*_*P*_ *× y*_*t−*1_), and allowed us to evaluate dynamic changes in the balance Δ_*S−P*_ = *β*_*S*_ *− β*_*P*_ between the two.

Consistent with recent findings in humans and mice^16,17^, Bayesian model comparison indicated a clear superiority of the two-state GLM-HMM over the standard one-state GLM in the S-ketamine experiment (*δ*_*BIC*_ = *−*3.65 *×* 10^3^). According to the two-state GLM-HMM, perception fluctuated between an internal mode, shaped by the stabilizing internal prediction *y*_*t−*1_, and an external mode, dominated by the SAR-weighted input *s*_*t*_. External mode increased Δ_*S−P*_ by 2.8 ± 0.29 (T(81) = 9.5, p = 5.22 *×* 10^−13^; Figure 2B-C). Switches between modes occurred in intervals of 179.97 ± 19.39 sec.

The presence of slow fluctuations between external and internal modes suggests that, instead of causing a constant increase in the sensitivity to external inputs, NMDAR hypofunction may affect perception by shifting the dynamic balance between the two modes. Indeed, S-ketamine did not alter the weights of the two-state GLM-HMM (Figure 2C), but increased the probability of external at the expense of internal mode (1.01 ± 0.03, z = 30.7, p = 4.26 *×* 10^−206^; Figure 2D). This effect was stable over time and present across the full range of SAR (Figure 2D). Inter-individual differences in the effects of S-ketamine confirmed that NMDAR hypofunction raised the sensitivity to sensory information (Figure 2A) by modulating the time participants spent in external and internal modes, respectively (*ρ* = 0.41, T(26) = 2.3, p = 0.03).

Strikingly, the data from the Scz-control study mirrored the effect of S-ketamine on the balance between external and internal mode: The two-state GLM-HMM outperformed the standard one-state GLM (patients: *δ*_*BIC*_ = *−*981.65; controls: *δ*_*BIC*_ = *−*862.91) and revealed two opposing modes (Δ_*S−P*_ = 1.44 ± 0.33, T(44) = 4.33, p = 3.39 *×* 10^−4^; Figure 2F) that alternated in intervals of 265.38 ± 57.76 sec for patients and 230.99 ± 65.04 sec for controls. Patients and controls did not differ with respect to the weights of the two-state GLM-HMM (Figure 2G). Instead, Scz patients spent more time in external mode (0.52 ± 0.03, z = 16.88, p = 1.23 *×* 10^−63^; Figure 2H).

We could not attribute between-mode transitions to fatigue, task difficulty (Figure 2D and H), executive function as reflected by response times (RT, Supplemental Figure S1C and S4), psychotomimetic effects of S-ketamine, global psychosis proneness, the clinical severity of Scz, stereodisparity thresholds, or subjective arousal, intoxication and nervousness (Supplemental Figure S5).

To our knowledge, these results are the first to uncover a neural mechanisms for the slow, task-related fluctuations in perceptual inference that have been observed across variety of tasks in humans and mice^14–17^. We found that healthy individuals who receive the NMDAR antagonist S-ketamine and patients diagnosed with Scz are prone to an external mode of perception. The external mode partly decouples perceptual inference from internal predictions that reflect prior knowledge about the world. In health, this may prevent circular inferences in recurrent neural networks, where predictive feedback modulates activity even at early stages of sensory processing^21,22^. A predominance of external mode, on the other hand, exposes perception to the destabilizing effects of ambiguity. Such transient failures of perceptual inference may cause individuals to be deluded by spurious connections between unrelated events^3^, to attribute the sensory consequences of their actions to an outside force, and to hallucinate signals in noise^3^.

During external mode, healthy participants were more confident in their choices (0.72 ± 0.07, z = 9.92, p = 7.85 *×* 10^−22^, Supplemental Figure S1E) and scored higher on the *Clinician-Administered-Dissociative-States-Scale*^23^ (CADSS, 1.05 ± 0.54, T(208.05) = 1.95, p = 0.05, Supplemental Figure S5B). Given the known association of psychosis with elevated confidence^24^ and dissociative symptoms^25^, intervals of external mode may thus reflect the computational correlate of individual psychotic experiences. The dynamic nature of between-mode transitions illustrates how constant and potentially heritable dysfunctions of the NMDAR, such as GRIN2A mutations in Scz^26^, may produce symptoms of psychosis that are recurrent and transient in nature.

## 3. Methods

### 3.1 Ressource availability

#### 3.1.1 Lead contact

Further information and requests for resources should be directed to and will be fulfilled by the lead contact, Veith Weilnhammer.

#### 3.1.2 Materials availability

This study did not generate new unique reagents.

#### 3.1.3 Data and code availability

All data and code associated with this study will be made available on the associated Github repository https://github.com/veithweilnhammer/modes_ketamine_scz upon publication. Key resources are listed in Supplemental Table S1.

### 3.2 S-ketamine vs. placebo

The S-ketamine experiment consisted in a total of three experimental sessions. During the first session, we screened participants for S-ketamine contraindications (arterial hypertension, prior psychiatric or neurological diagnoses including substance use disorder, use of psychoactive medication), and assessed psychosis proneness using the 40-item *Peters Delusion Inventory* (PDI^27^) and the 32-item *Cardiff Anomalous Perception Scale* (CAPS^28^). Moreover, we conducted three experimental pre-test runs that tested the ability to process stereodisparity (run 1, SAR = 1, cut-off: perceptual accuracy > 0.75), ensured the experience of spontaneous switches during bistable perception (run 2, SAR = 0, cut-off: perceptual stability < 0.96, corresponding to phase durations < 40 sec), and familiarized participants with the main experiment (run 3, see below for details).

In the subsequent two sessions, participants received a continuous intravenous infusion of either S-ketamine at 0.1 mg/kg/h or a saline placebo. Health screenings were repeated before each session to ensure the participants remained eligible. At each day of testing, we checked for alcohol intoxication using a breathalyzer and recent illicit substance use via a urine drug screen.

Our experimental protocol was double-blinded: The order of S-ketamine and placebo administration was counter-balanced across participants, with at least a two week interval between sessions. The participants, as well as the experimenters tasked with collecting the behavioral and psychometric data, were unaware of whether S-ketamine or placebo was administered by an independent group of clinicians who excluded undiagnosed psychotic illness using the *Brief Psychiatric Rating Scale* (BPRS^29^), established the intravenous line, started the infusion 15 min prior to the experiment, monitored the participants for side effects (blood pressure, drowsiness and vasovagal reactions, psychotomimetic effects), and removed the intravenous line at the end of the experiment, after which participants were monitored for at least 30 min. Deblinding occurred after data collection was complete.

#### 3.2.1 Sample characteristics

We screened a total of 87 right-handed individuals with (corrected-to-) normal vision, who were naive to the purpose of the study and gave written informed consent before participating. All experimental procedures were approved by the ethics committee at Charité Berlin.

From the group of screened participants, 31 did not meet our pretest criteria (6 due to perceptual accuracy < 0.75, 15 due to perceptual stability > 0.96, 8 due to substance use, 1 due to do a diagnosis of ADHD, and 1 due to medication with sertraline). Out of the remaining 56 participants who were eligible for the S-ketamine experiment, we aborted the main experiment in 1 participant due to high blood pressure at baseline (RR > 140/80 mmHG), in 2 participants due to strong psychotomimetic effects (micropsia) or dizziness under S-ketamine, and in 1 participant due to a vasovagal syncope during intravenous insertion. 24 participants were not available for the main experiment after successful pre-testing. We therefore report the data from a total of 28 participants (mean age: 28.93 ± 1.35 years, 18 female) who met all inclusion criteria and completed all experimental sessions.

#### 3.2.2 Experimental paradigm

We presented the experiment using Psychtoolbox 3^30^ running in Matlab R2021b (session 1: CRT-monitor at 85 Hz, 1280 x 1024 pixels, 60 cm viewing distance and 39.12 pixels per degree visual angle; session 2 and 32: CRT-monitor at 85Hz, 1280 x 1024 pixels, 40 cm viewing distance and 26.95 pixels per degree visual angle).

##### Procedure

Throughout the experiment, participants reported their perception of a discontinuous SMF stimulus (Supplemental Video S1). In this stimulus, random dots distributed on two intersecting rings induce the perception of a spherical object (diameter: 15.86°, rotational speed: 12 sec per rotation, rotations per block: 10, individual dot size: 0.12°) that rotates around a vertical axis with the front surface to the left or right^20^. Stimuli were presented in 120 sec blocks, separated by 10 sec fixation intervals.

Participants viewed the stimuli through a custom mirror stereoscope. In the pretest experiment, we presented stimuli at complete disambiguation (run 1, SAR = 1), full ambiguity (run 2, SAR = 0) and across five levels ranging from full ambiguity to complete disambiguation across five levels (run 3-5, *SAR* ∈ {0, 0.1, 0.25, 0.5, 1}). The SAR, which was constant within blocks, defines the fraction of stimulus dots that received a disambiguating 3D signal. Within each block, the direction of rotation enforced by the disambiguating 3D signal changes in average intervals of 10 sec (i.e., at a probability of 0.15 per stimulus overlap, see below). We pseudo-randomized the order of SAR across blocks and the direction of disambiguation within blocks.

Participants were naive to the potential ambiguity in the visual display, passively experienced the stimulus and reported changes in their perception alongside their confidence via button-presses on a standard USB keyboard (right middle-finger on d: rotation of the front-surface to the right at high confidence; right index-finger on f: rotation of the front-surface to the right at low confidence; left middle-finger on k: rotation of the front-surface to the left at high confidence; left index-finger on j: rotation of the front-surface to the left at low confidence; thumb on space bar: unclear direction of rotation). Unclear perceptual states occurred at a rate of 0.03 ± 0.01 and were excluded from further analyses.

The direction of rotation enforced by *s*_*t*_ (i.e., whether the parametric 3D signal enforced leftward or rightward rotation of the front surface) changed at a rate of 0.15 per overlap (i.e., on average every 10 sec). Changes in *s*_*t*_ and the order of blocks, each corresponding to one level of SAR, were pseudo-random.

In session 1 (pre-test), each run (runs 1 to 3) consisted of six blocks. In session 2 and 3 (main experiment), each run (run 4 and 5) consisted of 10 blocks. After every third block, the main experiment was paused to allow for the monitoring of the participants’ vital signs (blood pressure and pulse rate) and dynamic changes in psychotomimetic experiences. The latter was assessed using the 6 item *Clinician-Administered-Dissociative-States-Scale* (CADSS^23^) and three additional questions (Q1: *How awake do you feel?*, Q2: *How intoxicated do you feel?*, Q3: *How nervous do you feel?*) to which participants responded by clicking on a continuous line that encoded responses from *not at all* to *very much*. To measure global psychotomimetic effects of S-ketamine vs. placebo, participants completed the Questionnaire for the *Assessment of Altered States of Consciousness* (5D-ASC^31^) at the end of session 2 and 3. In addition, we collected responses on a debriefing questionnaire, in which we asked participants to describe whether they were able to accurately perceive the two directions of rotation induced by the SFM stimulus, whether they noticed any differences between blocks, whether they would guess that they received S-ketamine or placebo, and whether they had experienced any effects that they would attribute to a psychoactive substance.

##### Stereodisparity thresholds

At the beginning of the session 2 and 3, we conducted an independent stereo-acuity test to detect a potential effect of S-ketamine on stereodisparity thresholds^19^. We presented 5000 dots (each at 0.15° visual angle) within a square of 11° x 11° around a central fixation cross (0.10°). We added a stereodisparity signal to all dots on a Landolt C, i.e., a circle (1.37° radius, 2.06° width) with a 90° gap located either at the left, top, right or bottom. Stimuli were presented for 1 sec, after which participants reported the location of the gap by pressing the up-, down-, left- or right-arrow key within a 2 sec response interval, followed by 5 sec of fixation before the next trial.

We adjusted the stereodisparity of the Landolt C in a two-up-one-down staircase across 40 trials (initial stereodisparity: 0.0045°, correct response: decrease in the available stereodisparity by one step; incorrect response: increase by two steps, initial step-size: 0.001°, reduction to 0.0005° after first reversal). Stereodisparity thresholds were defined by the average stereodisparity present at the last 10 trials of the staircase.

##### Scores and Questionnaires

Supplementary Table S2 provides an overview of our psychometric data.

### 3.3 Scz patients vs. healthy controls

To test whether Scz patients show similar changes in bimodal inference as healthy participants who receive the NMDAR-antagonist S-ketamine, we re-analyzed data from a previously published case-control study^19^ that compared Scz patients to healthy participants in paradigm analogous to the S-ketamine experiment described above.

#### 3.3.1 Sample characteristics

We report data from 23 patients diagnosed with paranoid Scz (ICD-10: F20.0, 18 male, age = 37.13±2.42) and 23 controls (17 male, age = 33.57±1.74) that were matched for gender, age and handedness^19^.

#### 3.3.2 Experimental paradigm

Stimuli were presented using Psychtoolbox 3^30^ running in Matlab R2007b (CRT-Monitor at 60 Hz, 1042x768 pixels, 59.50cm viewing distance, 30.28 pixels per degree visual angle).

##### Main Experiment

Throughout the experiment, participants reported their perception of a discontinuous SFM stimulus (see Supplemental Video S2) via button-presses on a standard USB keyboard. In contrast to the S-ketamine experiment, the 300 dots (0.05°) that composed the stimulus (2.05°x 2.05°) were not placed on rings, but on a Lissajous band defined by the perpendicular intersection of two sinusoids (*x*(*p*) = *sin*(*A* * *p*) and *y*(*p*) = *cos*(*B* * *p* + *δ*) with *A* = 3, *B* = 6, with *δ* increasing from 0 to 2*π* at 0.15 Hz. Overlapping configurations of the stimulus occurred in intervals of 3.33 sec. Participants viewed the stimuli through a mirror stereoscope. Fusion was supported by rectangular fusion-frames and a background of random dot noise (700 dots of 0.05° which moved at a speed of 1.98° per sec and changed their direction at a rate of 1 Hz).

We presented participants with 3 sessions of the main experiment, each consisting of 14 40.08 sec blocks that were separated by 5 sec of fixation and differed with respect to the SAR, ranging from full ambiguity to complete disambiguation in 8 levels (*SAR* ∈ {0, 0.01, 0.04, 0.9, 0.16, 0.26, 0.50, 1}). The frequency of changes in the direction of the disambiguating signal corresponded to the frequency of spontaneous changes that participants perceived during full ambiguity^19^ (SAR = 0). In contrast to the S-ketamine experiment, participants only reported the perceived direction of rotation *y*_*t*_ (left vs. rightward movement of the front surface).

##### Stereodisparity thresholds

We assessed stereodisparity thresholds in Scz patients and controls using the procedure described above.

##### Scores and Questionnaires

We used the PDI^27^ and the CAPS^28^ to measure delusional ideation and perceptual anomalies in Scz patients and controls. Clinical symptom severity was assessed using the *Positive and Negative Syndrome Scale* (PANSS)^32^.

### 3.4 Quantification and statistical procedures

This manuscript was written in RMarkdown. All data and summary statistics can be reviewed by cloning the Github respository https://github.com/veithweilnhammer/Ketamin_RDK and running the file *ketamine_scz_frmri_modes*.*Rmd*, which will be made public at the time of publication.

The SFM stimuli used in the above studies share an important feature: Even though physically ambiguous at all angles of rotation, spontaneous changes in the perceived direction of rotation are limited to overlapping configurations of the stimuli^19,20^ (see also Supplemental Figure S1 and S4). This is because depth-symmetry, which is a prerequisite for changes in subjective experiences during bistable SFM^19,20^, is limited to timepoints when the bands that compose the stimuli overlap (Supplemental Video S1 and S2).

We therefore discretized the perceptual timecourse of all experiments into a sequence of overlaps that occur at times *t* (1.5 sec inter-overlap interval for the S-ketamine and fMRI experiment, 3.33 sec inter-overlap interval for the case-control experiment). Each inter-overlap interval is characterized by the primary independent variable *s*_*t*_ = [*−*1, 1] *× SAR* (the SAR-weighted input ranging from maximum information for leftward rotation to maximum information for rightward rotation). As secondary independent variables, we considered block and session index (reflecting the time participants were exposed to the experiment), participant identifiers and, if applicable, treatment or group identifiers. Primary dependent variables were *y*_*t*_ = [0, 1] (the experience of either leftward or rightward rotation), *r*_*t*_ (the time between the button-press indicating a perceptual event relative to the preceding overlap) and, if applicable, *c*_*t*_ = [0, 1] (low vs. high confidence). As secondary dependent variables, we computed perceptual accuracy (the probability of *y*_*t*_ *≅ s*_*t*_) and perceptual stability (the probability of *y*_*t*_ = *y*(*t −* 1)). We report averages as mean ± s.e.m.

#### 3.4.1 Conventional statistics

The goal of our conventional statistics was to quantify the effect of NMDAR hypofunction, whether due to an antagonism with S-ketamine or due to a diagnosis of Scz, on the interpretation of ambiguous sensory information. To this end, we performed standard logistic and linear regression by fitting (general) linear effects models using the R-packages lmer, glmer and afex (see Supplemental Table S2). We predicted *y*_*t*_, *c*_*t*_, perceptual accuracy and perceptual stability in logistic regression, and *r*_*t*_ in linear regression. We estimated random intercepts defined within participants in the S-ketamine experiment and nested random intercepts for participants within groups in the case-control experiment. We applied a Bonferroni-correction for the number of main effects and interactions within models. For non-normally distributed secondary dependent variables, we performed rank-based tests to assess correlations (Spearman) and distribution differences (Wilcoxon).

#### 3.4.2 Computational modeling

Having established the effect of NMDAR hypofunction on the interpretation of ambiguous sensory information, we used computational modeling to arbitrate between two mechanistic explanations on how S-ketamine and schizophrenia may alter perceptual inference.

##### Hypothesis H1

**Unimodal inference** In one scenario, NMDAR hypofunction may induce a global increase in the sensitivity to external inputs relative to the stabilizing internal prediction. This unimodal scenario would be reflected by S-ketamine- or Scz-related changes in the weights *w* ≡ {*β*_*S*_, *β*_*P*_, *β*_*B*_} of a GLM that predicts percepts *y*_*t*_ from the input vector *x*_*t*_, which consists in the SAR-weighted external input *s*_*t*_, the stabilizing internal prediction *y*_*t−*1_ and a constant bias *b*:

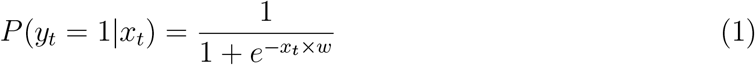

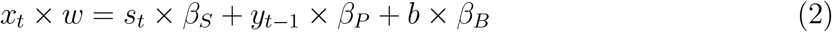

According to the unimodal hypothesis 1, NMDAR hypofunction increases *β*_*S*_ at the expense of *β*_*P*_, leading to an increase of Δ_*S−P*_ = *β*_*S*_ *− β*_*P*_ .

##### Hypothesis H2

**Bimodal inference** In an alternative scenario, NMDAR hypofunction does not change the weights of the GLM directly, but modulates the transition between latent modes^17^ or decision-making strategies^16^ that differ with respect to the balance between external inputs *s*_*t*_ and the stabilizing internal prediction provided by *y*_*t−*1_. In the bimodal scenario, perceptual inference is characterized by two latent modes *z*_*t*_ (i.e., states in a HMM) that alternate at a probability per overlap that is defined by a 2 x 2 transition matrix *A*:

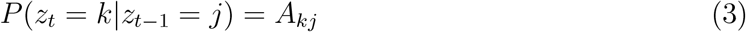

Each state *z*_*t*_ is associated by an independent GLM defined by the weights *w*_*k:*_

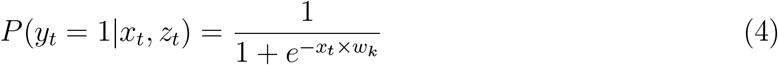

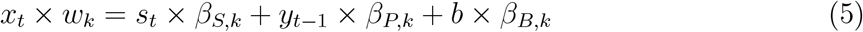

The bimodal hypothesis H2 differs from the unimodal hypothesis H1 in two ways: First, bimodal inference is characterized by two (as opposed to one) GLMs that differ with respect to Δ_*S−P*_ : During external mode, *β*_*S*_ is increased relative to *β*_*P*_, whereas during internal mode, *β*_*P*_ is increased relative to *β*_*S*_. Second, during bimodal inference, NMDAR hypofunction does not alter the weights within the external and internal GLMs, but modulates the transition probability between the two.

##### Procedure

To contrast hypotheses H1 and H2, we fitted unimodal and bimodal GLM-HMMs using SSM^33^ (Supplemental Table S2), compared models via Bayesian Information Criterion (BIC), and assessed the effects of S-ketamine or Scz on the posterior model parameters, i.e., HMM transition probabilities and the mode-dependent GLM weights *w*_*k*_. Model fitting using SSM is governed by the hyperparameters *σ*^2^ and *α. σ*^2^ denotes the variance of a prior over the GLM weights *w*_*k*_. Smaller values of *σ*^2^ shrink *w*_*k*_ toward 0, whereas *σ* = ∞ leads to flat priors. We set *σ*^2^ to 100 for GLMs that predicted group-level data, and to 1 for GLMs that predicted participant- or session-level data, which were initialized with group-level estimates of *w*_*k*_. *α* defines the Dirichlet prior over the transition matrix *A* and is flat for *α* = 1. We set *α* to 1 for all group-level and participant-level fits.

For each experiment, computational modeling was carried out in a sequence of 3 steps: In a first step, we fitted a unimodal GLM initialized with noisy weights to the group-level data (i.e., data pooled across participants within an individual experiment) for a total of n = 100 iterations and computed the average posterior weights *w*_*n*_. In a second step, we fitted the group-level data with the unimodal and the bimodal GLM-HMM initialized by *w*_*n*_, extracted the posterior parameters *w*_*k*_, and compared the models using BIC.

In a third step, we fitted the unimodal and the bimodal GLM-HMM to session-level data (S-ketamine experiment) and participant-level data (case-control experiment). Models were initialized by the average weights *w*_*n*_ of the corresponding group-level model. For all bimodal group-, participant- and session-level GLM-HMMs, we defined the latent mode associated with the higher posterior *β*_*S*_ estimate as external. Our definition of mode is thus agnostic with respect to *β*_*P*_ and Δ_*S−P*_. This allowed us to contrast external-to-internal bimodal inference (hypothesis H2) with a third alternative where perception fluctuates between latent states that differ with respect to decision noise (**hypothesis H0**). Low decision noise is characterized by higher posterior estimates of *β*_*S*_ and *β*_*P*_, whereas high decision noise is characterized by lower posterior estimates of *β*_*S*_ and *β*_*P*_. Δ_*S−P*_ therefore discerns fluctuations in decision noise (H0, no changes in Δ_*S−P*_) from external-to-internal bimodal inference (H2, mode-associated changes in Δ_*S−P*_ with higher estimates during external mode).

For summary statistics, we extracted the posterior weights *w*_*k*_ (separately for external and internal mode) and the dynamic posterior probability of external mode *z*_*t*_ = *e*.

## 5 Supplemental Information

### 5.1 Supplemental Figure S1

**Supplemental Figure S1.**
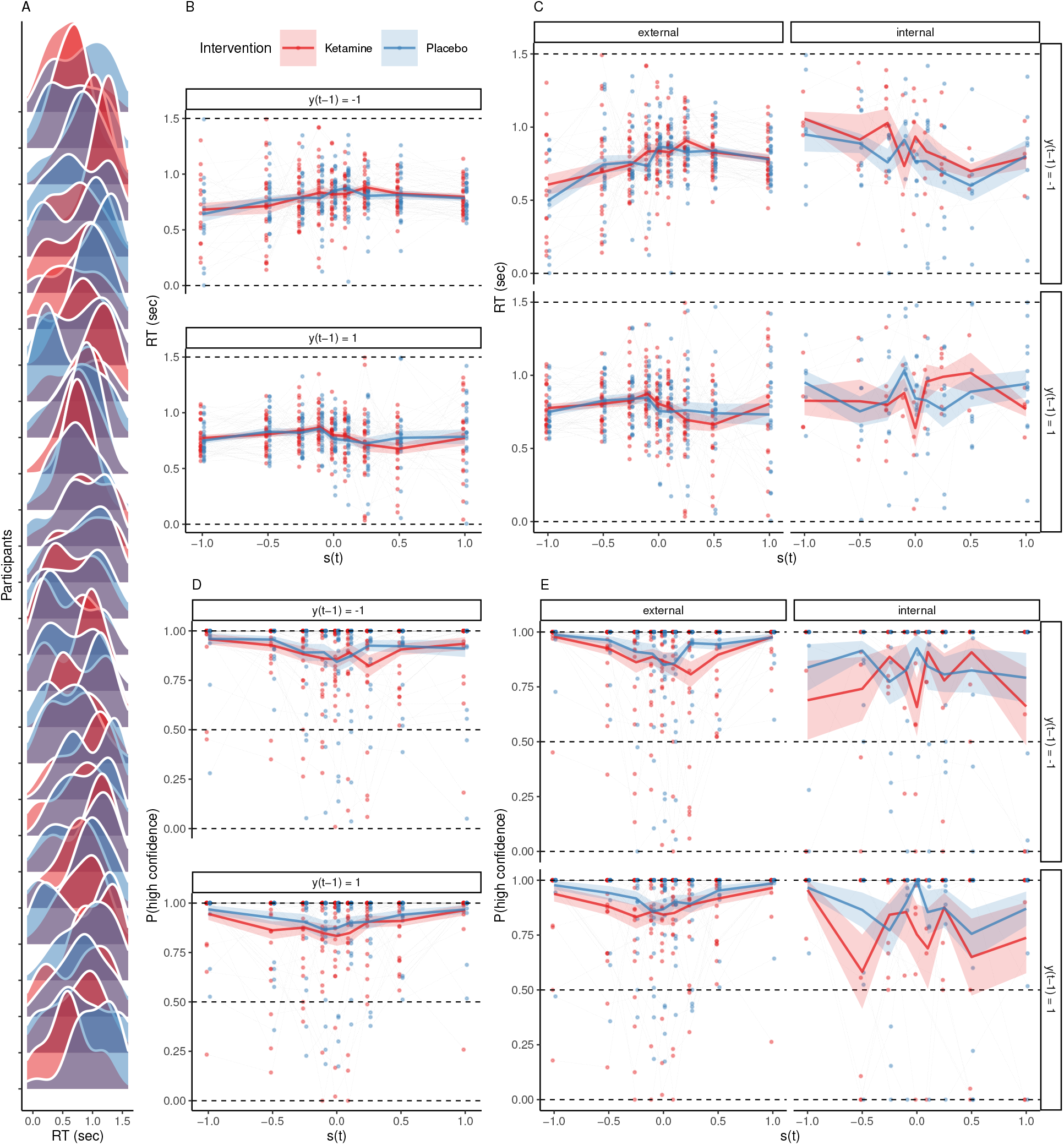
The effects of ketamine and bimodal inference on RT. **A**.RT were non-uniformly distributed across the inter-overlap interval (D = 0.09, p = 5.38 *×* 10^−9^, one-sample Kolmogorov-Smirnov test). This corroborates that changes in perception aligned with the overlapping configurations of the stimulus after S-ketamine (red) and placebo (blue). **B**.RT showed a quadratic relationship with *s*_*t*_ (*−*6.87 ± 1.68, T(6.2 *×* 10^3^) = *−*4.1, p = 5.1 *×* 10^−4^), indicating faster responses when sensory information was reliable (|*s*_*t*_| ≫ 0; note that SAR as shown in Figure 2A and 2E is equal to |*s*_*t*_|). We observed no main effect of S-ketamine (red) vs. placebo (blue) on RT (*−*3.35 *×* 10^−3^ ± 0.01, T(6.2 *×* 10^3^) = *−*0.32, p = 1). **C**.We found no additional effect of mode on RT (0.02 ± 0.03, z = 5.96 *×* 10^3^, p = 0.78). **D**.Confidence showed a quadratic relationship with *s*_*t*_ (74.83 ± 2.39, z = 31.32, p = 3.22 *×* 10^−214^), confirming that participants were more confident when sensory information was reliable (|*s*_*t*_| = *SAR* ≫ 0). Relative to placebo (blue), S-ketamine (red) reduced choice confidence (*−*0.21 ± 0.04, z = *−*5.9, p = 4.36 *×* 10^−8^), and decreased the quadratic effect of *s*_*t*_ on confidence (*−*19.95 ± 2.36, z = *−*8.45, p = 3.48 *×* 10^−16^). **E**.External mode increased confidence globally (0.72 ± 0.07, z = 9.92, p = 7.85*×*10^−22^) and by elevating the quadratic effect of *s*_*t*_ on confidence (242.61 ± 18.43, z = 13.16, p = 3.37 *×* 10^−38^). When controlling for mode, the negative effect of S-ketamine (red) vs. placebo (blue) on confidence and on the quadratic relationship of confidence with *s*_*t*_ remained significant.

### 5.2 Supplemental Figure S2

**Supplemental Figure S2.**
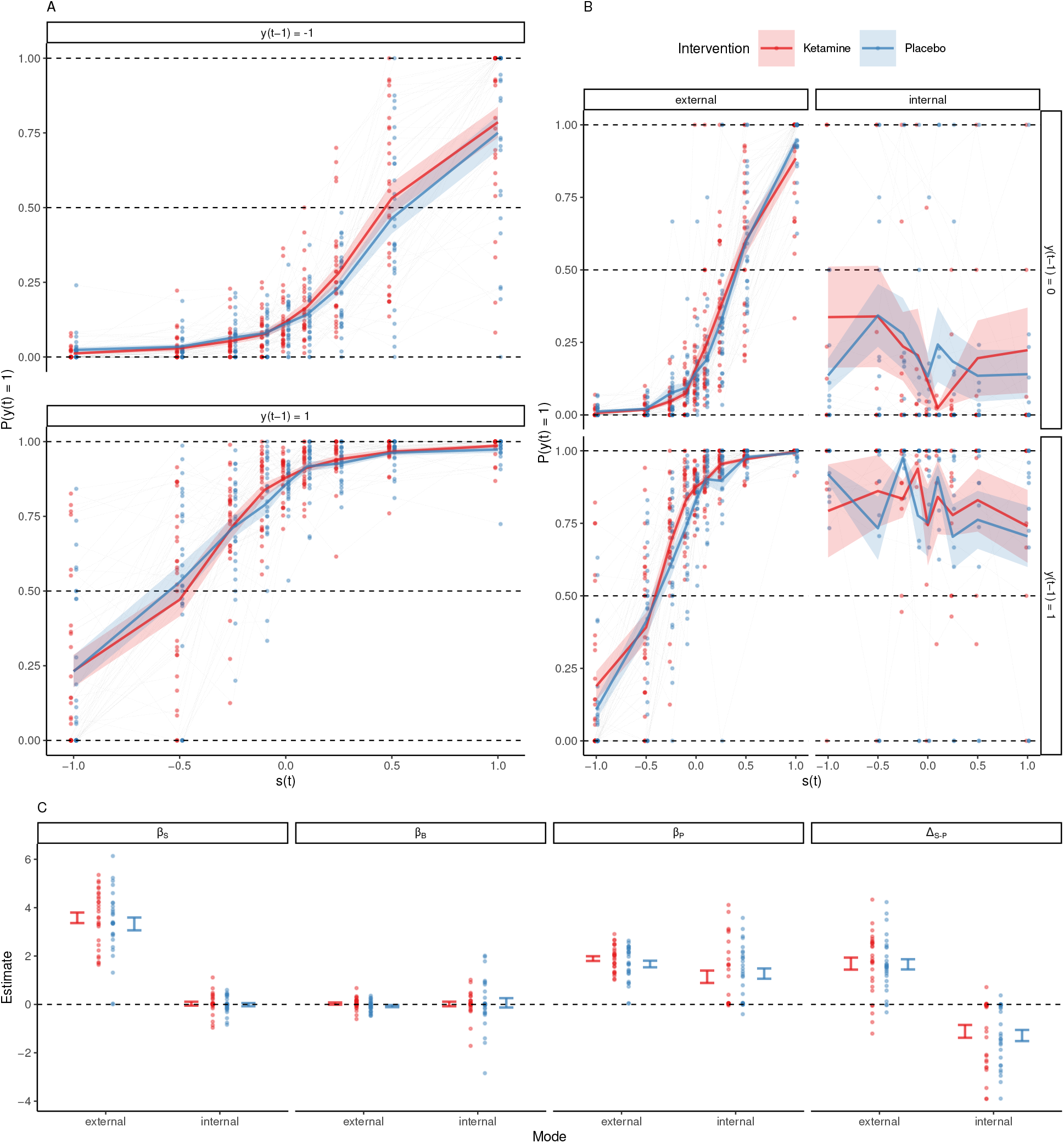
Extended data on the effects of S-ketamine and mode on perceptual inference (related to Figure 2A-C). **A**.Here, we show psychometric curves (percept *y*_*t*_ versus input *s*_*t*_) under S-ketamine (red) and placebo (blue). The plot separates times *t* for which the previous experience was leftward rotation (*y*_*t−*1_ = *−*1, upper panel) and rightward rotation (*y*_*t−*1_ = +1, lower panel). As expected, *y*_*t*_ was driven by both the external input *s*_*t*_ (*β*_*S*_ = 3.01 ± 0.06, z = 50.39, p < 2.2 *×* 10^−308^) and the previous percept *y*_*t−*1_ (*β*_*P*_ = 2.06 ± 0.03, z = 80.58, p < 2.2 *×* 10^−308^). We found no significant interaction between the *s*_*t*_ and *y*_*t−*1_ (*−*0.06 ± 0.06, z = *−*1.06, p = 1). Relative to placebo, S-ketamine caused a shift of *y*_*t*_ toward *s*_*t*_ (0.45 ± 0.08, z = 5.6, p = 1.71 *×* 10^−7^), with no significant effect on *y*_*t−*1_ (0.08 ± 0.04, z = 2.39, p = 0.13). We found no significant three-way-interaction (drug x *s*_*t*_ x *y*_*t−*1_, *−*0.07 ± 0.08, z = *−*0.9, p = 1). **B**.This panel shows the data from panel (A) separately for times *t* where the HMM identified the mode of perceptual inference as external (left panels) or internal (right panels). When the mode of perceptual processing was added to the prediction of *y*_*t*_ from *s*_*t*_ and *y*_*t−*1_, the effect S-ketamine (red) vs. placebo (blue) on *s*_*t*_ disappeared (0.24 ± 0.11, z = 2.13, p = 0.53). Instead, changes in the balance between *s*_*t*_ and *y*_*t−*1_ were loaded onto fluctuations between external and internal mode, which caused perception to shift away from external inputs *s*_*t*_ (*−*4.23 ± 0.21, z = *−*20.01, p = 7.54 *×* 10^−88^) and toward previous experiences *yt −* 1 (0.78 ± 0.09, z = 8.64, p = 8.81 *×* 10^−17^). **C**.Here, we plot the weights from the GLM *y*_*t*_ = *β*_*S*_ *× s*_*t*_ + *β*_*P*_ *× y*_*t−*1_ + *β*_*B*_ *×* 1, alongside the balance between external inputs and previous experiences Δ_*S−P*_ = *β*_*S*_ *− β*_*P*_ during external and internal mode. Colors indicate S-ketamine (red) and placebo (blue). *β*_*S*_, the weight associated with the external input *s*_*t*_, was positive in external mode, but reduced to zero in internal mode (*−*3.55 ± 0.23, T(81) = *−*15.44, p = 4.78 *×* 10^−24^). We found no additional effect of S-ketamine (red) versus placebo (blue; *−*0.25 ± 0.23, T(81) = *−*1.1, p = 1) and no significant interaction (0.21 ± 0.33, T(81) = 0.65, p = 1). *β*_*B*_, the weight associated with the constant response bias *b* toward rightward rotation, was not different from zero (*β*_*B*_ = 0.04 ± 0.11, T(98.36) = 0.31, p = 1). We found no effect of drug (*−*0.11 ± 0.14, T(81) = *−*0.74, p = 1) or mode (*−*0.02 ± 0.14, T(81) = *−*0.12, p = 1) on the bias weight *β*_*B*_. *β*_*P*_, the weight associated with the previous percept *y*_*t−*1_ was not modulated by S-ketamine (*−*0.22 ± 0.26, T(81) = *−*0.87, p = 1) or mode (*−*0.75 ± 0.26, T(81) = *−*2.92, p = 0.29). There was no significant interaction between drug and mode with respect to *β*_*P*_ (0.35 ± 0.36, T(81) = 0.97, p = 1). The balance Δ_*S−P*_ between external inputs and internal predictions was determined by mode (2.8 ± 0.29, T(81) = 9.5, p = 5.22 *×* 10^−13^), with no significant effect of S-ketamine (0.03 ± 0.29, T(81) = 0.1, p = 1) and no interaction (0.14 ± 0.42, T(81) = 0.34, p = 1).

### 5.3 Supplemental Figure S3

**Supplemental Figure S3.**
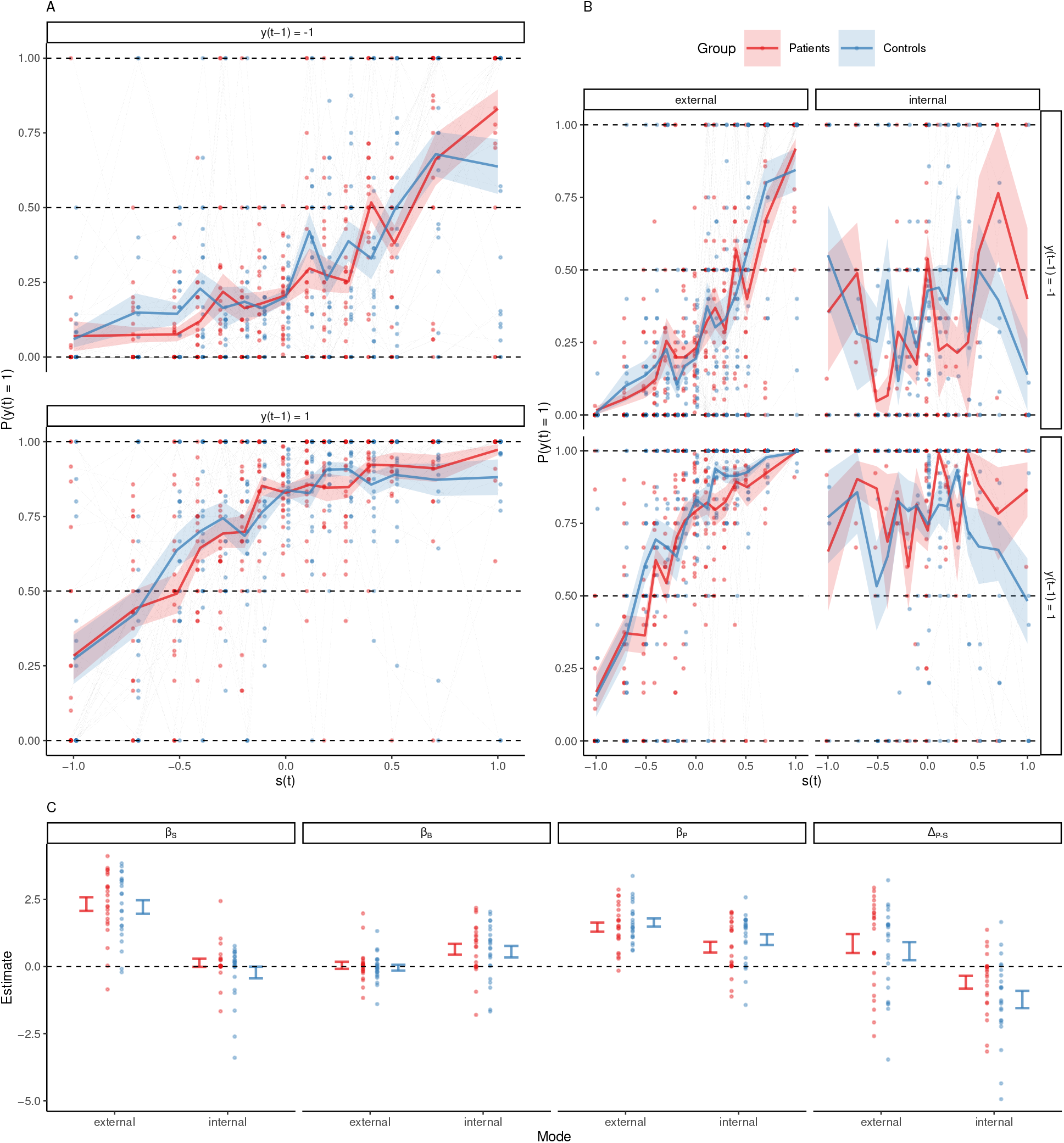
Extended data on external and internal mode in Scz patients and healthy controls (related to Figure 2E-H). **A**.Here, we show psychometric curves (percept *y*_*t*_ versus input *s*_*t*_) in patients (red) and controls (blue). The plot separates times *t* for which the previous experience was leftward rotation (*y*_*t−*1_ = *−*1, upper panel) and rightward rotation (*y*_*t−*1_ = +1, lower panel). Perception was driven by *s*_*t*_ (*β*_*S*_ = 2.77 ± 0.11, z = 24.85, p = 2.18 *×* 10^−135^) and *y*_*t−*1_ (*β*_*P*_ = 1.5 ± 0.03, z = 58.2, p < 2.2 *×* 10^−308^), with no significant interaction between *s*_*t*_ and *y*_*t−*1_ (*−*5.41 *×* 10^−3^ ± 0.11, z = *−*0.05, p = 1). Patients were more sensitive to *s*_*t*_ (0.75 ± 0.15, z = 4.96, p = 5.6 *×* 10^−6^). We found no significant three-way-interaction (group x *s*_*t*_ x *y*_*t−*1_, *−*0.37 ± 0.15, z = *−*2.45, p = 0.11). **B**.This panel shows the data from panel (A) separately for times *t* where the HMM identified the mode of perceptual inference as external (left panels) or internal (right panels). When the mode of perceptual processing was added to the prediction of *y*_*t*_ from *s*_*t*_ and *y*_*t−*1_, the difference between patients (red) and controls (blue) in the effect of *s*_*t*_ on *y*_*t*_ disappeared (*−*0.02 ± 0.22, z = *−*0.08, p = 1). Instead, changes in the balance between *s*_*t*_ and *y*_*t−*1_ were loaded onto fluctuations between external and internal mode, which caused perception to shift away from external inputs *s*_*t*_ (*−*3.47 ± 0.29, z = *−*11.95, p = 1.01 *×* 10^−31^) and toward previous experiences *yt −* 1 (0.5 ± 0.07, z = 6.85, p = 1.15 *×* 10^−10^). **C**.Here, we plot the weights from the GLM *y*_*t*_ = *β*_*S*_ *× s*_*t*_ + *β*_*P*_ *× y*_*t−*1_ + *β*_*B*_ *×* 1, alongside the balance between external inputs and previous experiences Δ_*S−P*_ = *β*_*S*_ *− β*_*P*_ during external and internal mode. Colors indicate the group (patients in red, controls in blue). *β*_*S*_, the weight associated with the external input *s*_*t*_, was positive in external mode, but reduced to zero in internal mode (*−*2.19 ± 0.24, T(44) = *−*9.13, p = 4.07 *×* 10^−11^). We found no additional effect of group (*−*0.11 ± 0.37, T(87.69) = *−*0.3, p = 1) and no significant interaction (*−*0.25 ± 0.34, T(44) = *−*0.74, p = 1). *β*_*B*_, the weight associated with the constant response bias *b* toward rightward rotation, was not different from zero (0.05 ± 0.18, T(1.62 *×* 10^−8^) = 0.29, p = 1). We found no effect of group (*−*0.09 ± 0.25, T(1.62 *×* 10^−8^) = *−*0.37, p = 1). There was a trend for a positive effect of internal mode (0.6 ± 0.24, T(88) = 2.47, p = 0.06) on the bias weight *β*_*B*_. *β*_*P*_, the weight associated with the previous percept *y*_*t−*1_, was reduced in internal mode (*−*0.75 ± 0.26, T(88) = *−*2.92, p = 0.02), but not modulated by group (0.17 ± 0.32, T(9.88 *×* 10^−10^) = 0.54, p = 1). There was no significant interaction between group and mode with respect to *β*_*P*_ (0.11 ± 0.36, T(88) = 0.3, p = 1). The balance Δ_*S−P*_ between external inputs and internal predictions was determined by mode (1.44 ± 0.33, T(81) = 9.5, p = 3.39 *×* 10^−4^), with no significant effect of group (0.28 ± 0.54, T(87.97) = 0.52, p = 1) and no interaction (0.36 ± 0.47, T(44) = 0.76, p = 1).

### 5.4 Supplemental Figure S4

**Supplemental Figure S4.**
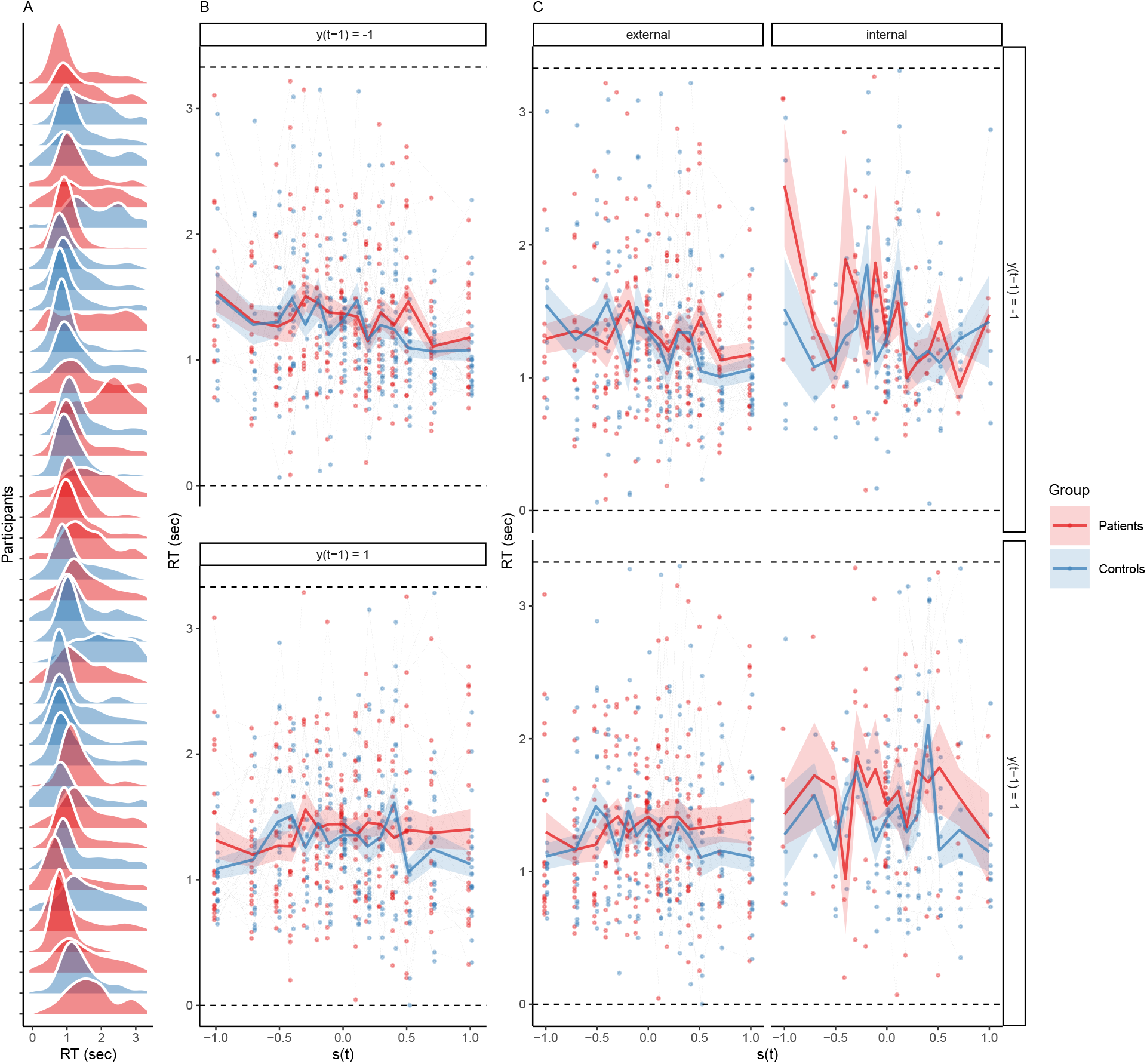
RT and bimodal inference in Scz patients and controls. **A**.RT were non-uniformly distributed across the inter-overlap interval (D = 0.22, p < 2.2 *×* 10^−308^, one-sample Kolmogorov-Smirnov test against uniformity) in patients (red) and controls (blue). This confirmed that changes in perception were aligned with the overlapping configurations of the stimulus. **B**.RT did not differ between patients (red) and controls (blue; *−*0.07 ± 0.08, T(66.96) = *−*0.87, p = 1). We found no quadratic relationship between RT and *s*_*t*_ (*−*3.54 ± 2.34, T(5.33 *×* 10^3^) = *−*1.51, p = 1). **C**.We found no effect of mode on RT (0.03 ± 0.04, z = 4.89 *×* 10^3^, p = 0.76).

### 5.5 Supplemental Figure S5

**Supplemental Figure S5.**
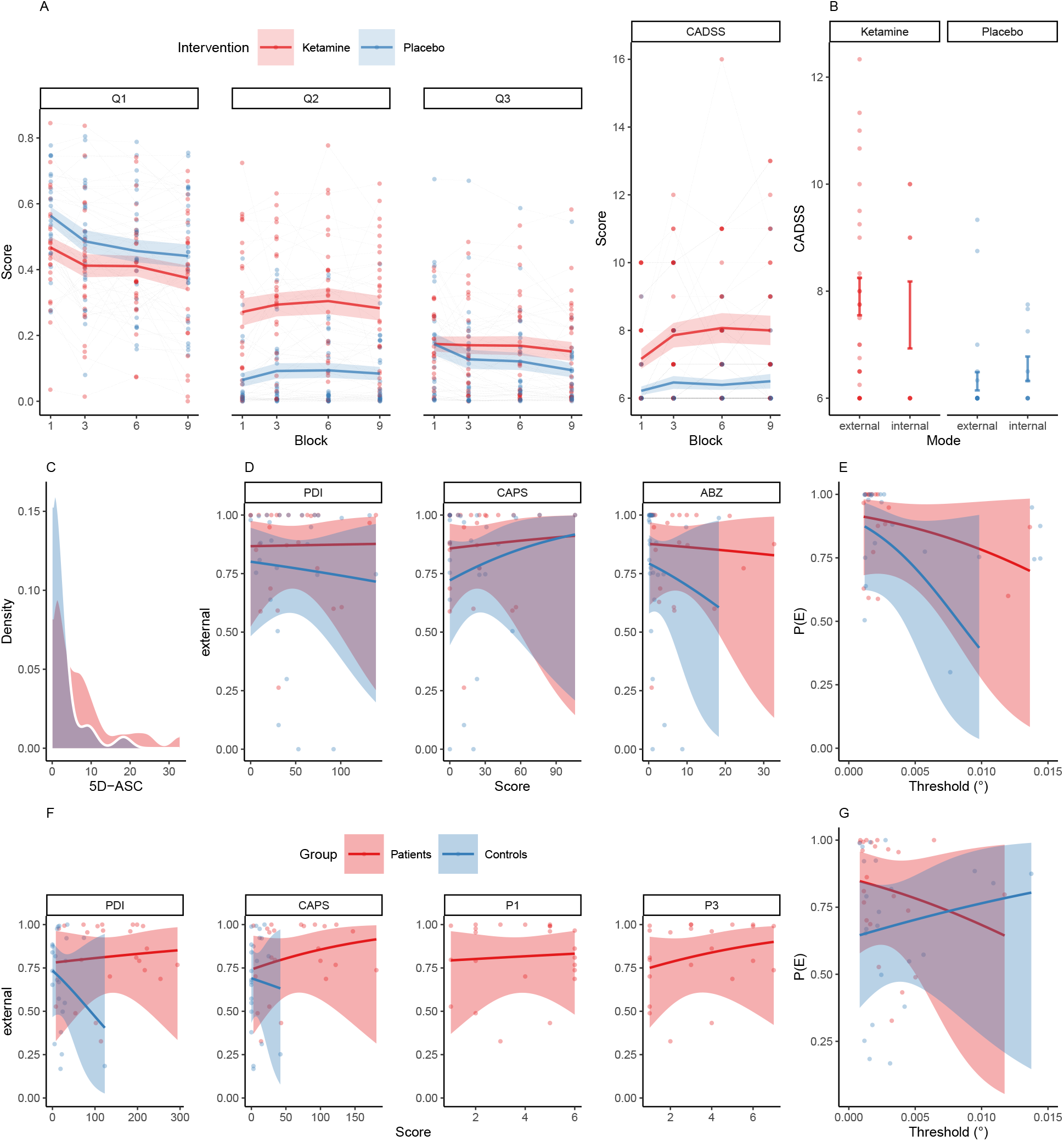
Scores and Questionnaires. **A**. Responses to Q1 (*How awake do you feel?*) indicated that participants felt more tired under S-ketamine (red) than placebo (blue; *−*1.53 ± 0.6, z = *−*2.57, p = 0.04), with no significant effect of time or a between-factor interaction. Responses to Q2 (*How intoxicated do you feel?*) indicated that participants felt more intoxicated under S-ketamine (3.32 ± 1.44, z = 2.3, p = 0.09), with no significant effect of time or a between-factor interaction. Responses to Q3 (*How nervous do you feel?*) revealed no effect of S-ketamine (*−*3.01 ± 2.62, z = *−*1.15, p = 1), time, nor a significant between-factor interaction. CADSS scores were elevated under S-ketamine (1.01 ± 0.34, T(185.32) = 2.99, p = 0.01) with a borderline trend for an increase over time (0.09 ± 0.04, T(185.61) = 2.24, p = 0.1) and no significant between-factor interaction. **B**. Q1-3 and CADSS scores were collected after blocks 1, 3, 6 and 9. To assess how the mode of perceptual inference was linked to dissociative symptoms, we separated the participants ratings according to the mode that dominated perception at the very end of the preceding block. While controlling the effect of S-ketamine (red) vs placebo (blue), we found that external mode increased dissociative symptoms (1.05 ± 0.54, T(208.05) = 1.95, p = 0.05), but had no effect on wakefulness (Q1), subjective intoxication (Q2) or nervousness (Q3). **C**. 5-ASC scores were elevated under S-ketamine (red) relative to placebo (blue; 4.89 ± 1.59, T(27.14) = 3.08, p = 9.33 *×* 10^−3^). **D**. Neither PDI, CAPS, nor 5-ASC scores were predictive of the probability of external mode (shown separately for S-ketamine in red and placebo in blue). **E**. Stereodisparity thresholds were not predictive of the probability of external mode (*−*28.73 ± 781.1, z = *−*0.04, p = 0.97). Thresholds did not differ between S-ketamine (red) and placebo (blue; W = 102, p = 0.66). **F**. Neither PDI, CAPS (patients in red and controls in blue), nor the PANSS items P1 (delusions) or P3 (hallucinations, patients only) predicted the probability of external mode. **G**.In patients (red) and controls (blue), stereodisparity thresholds were not predictive of the probability of external mode (*−*1.88 ± 2.05, z = *−*0.92, p = 1). Thresholds did not differ between groups (V = 976, p = 0.52).

### 5.6 Supplemental Table S1

**Supplemental Table S1.**
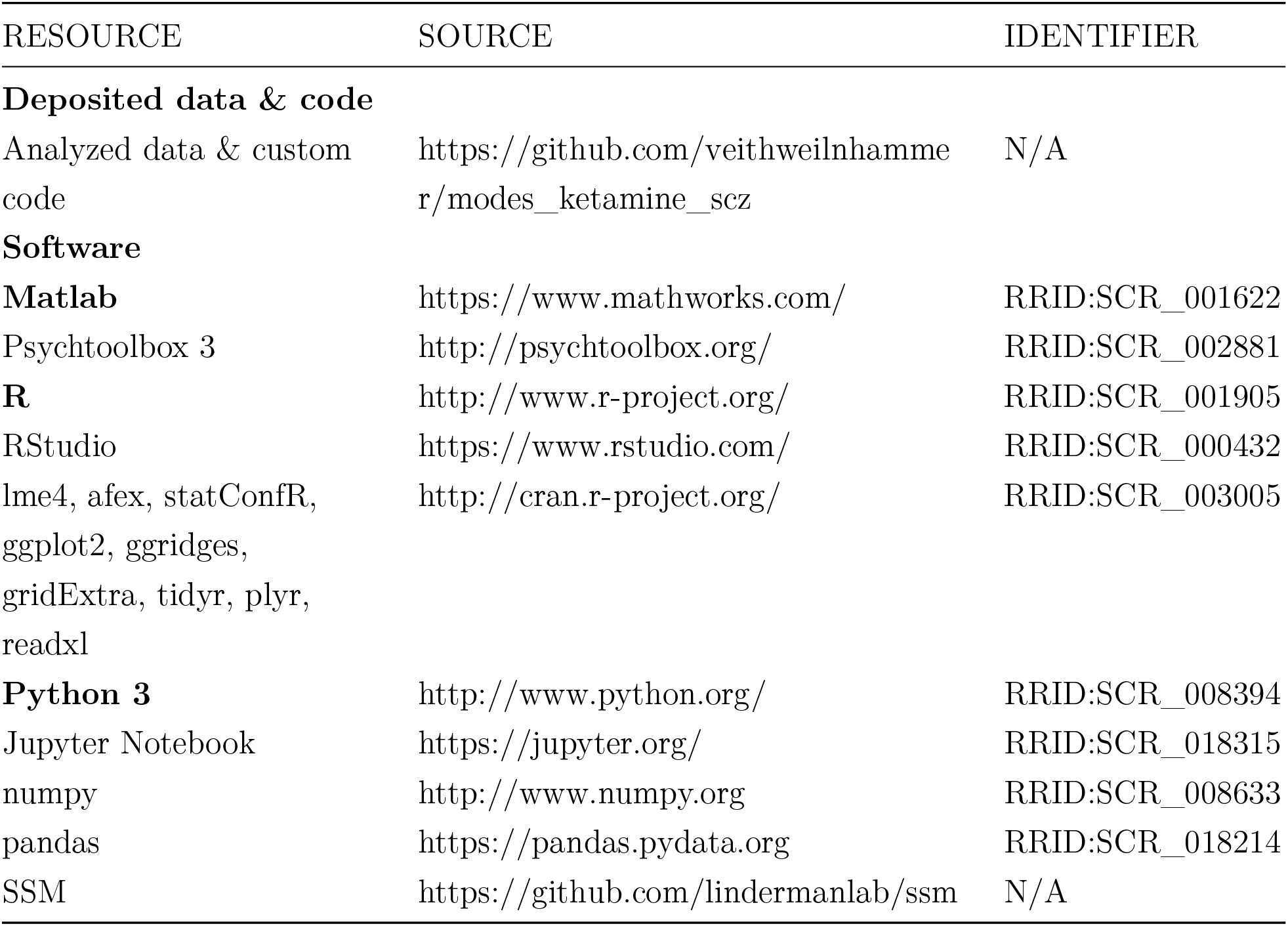
Key resources.

### 5.7 Supplemental Table S2

**Supplemental Table S2.** Psychometric data for the S-ketamine experiment.

| Scale | Scope | Condition | mean ± s.e.m. |
| --- | --- | --- | --- |
| **PDI**<sup>27</sup> | Delusion proneness | Global | $46.22 \pm 7.19$ |
| **CAPS**<sup>28</sup> | Hallucination proneness | Global | $23 \pm 5.05$ |
| **BPRS**<sup>29</sup> | Screen for psychotic illness | Global | $0.64 \pm 0.27$ |
| **5D-ASC**<sup>31</sup> | Altered states of consciousness | S-ketamine | $7.11 \pm 1.59$ |
| | | Placebo | $2.2 \pm 0.75$ |
| **CADSS**<sup>23</sup> | Dissociation | S-ketamine | $7.8 \pm 0.33$ |
| | | Placebo | $6.43 \pm 0.17$ |
| **Q1** | Wakefulness | S-ketamine | $0.41 \pm 0.03$ |
| | | Placebo | $0.48 \pm 0.03$ |
| **Q2** | Intoxication | S-ketamine | $0.29 \pm 0.03$ |
| | | Placebo | $0.09 \pm 0.02$ |
| **Q3** | Nervousness | S-ketamine | $0.17 \pm 0.02$ |
| | | Placebo | $0.13 \pm 0.03$ |
| **Stereovision** | Disparity thresholds | S-ketamine | $2.89 \times 10^{-3} \pm$ |
| | | | $6.18 \times 10^{-4}$ |
| | | Placebo | $2.75 \times 10^{-3} \pm$ |
| | | | $4.39 \times 10^{-4}$ |

### 5.8 Supplemental Table S3

**Supplemental Table S3.** Psychometric data for Scz-control-study.

| Scale | Scope | Condition | mean ± s.e.m. |
| --- | --- | --- | --- |
| **PDI**<sup>27</sup> | Delusion proneness | Patients | 138.83 ± 16.64 |
| | | Controls | 21.87 ± 5.75 |
| **CAPS**<sup>28</sup> | Hallucination proneness | Patients | 65.17 ± 10.56 |
| | | Controls | 7.13 ± 2.2 |
| **P1** | Delusions | Patients | 3.83 ± 0.39 |
| **P3** | Delusions | Patients | 3.35 ± 0.44 |
| **Stereovision** | Disparity thresholds | Patients | 2.82 $\times 10^{-3} \pm$ |
| | | | 5.13 $\times 10^{-4}$ |
| | | Controls | 3.46 $\times 10^{-3} \pm$ |
| | | | 7.14 $\times 10^{-4}$ |

